# Coordinated differentiation of human intestinal organoids with functional enteric neurons and vasculature

**DOI:** 10.1101/2023.11.06.565830

**Authors:** Charlie J. Childs, Holly M. Poling, Kevin Chen, Yu-Hwai Tsai, Angeline Wu, Caden W. Sweet, Abigail Vallie, Madeline K. Eiken, Sha Huang, Ryan Schreiner, Zhiwei Xiao, Ansley S. Conchola, Meghan F. Anderman, Emily M. Holloway, Akaljot Singh, Roman Giger, Maxime M. Mahe, Katherine D. Walton, Claudia Loebel, Michael A. Helmrath, Shahin Rafii, Jason R. Spence

## Abstract

Human intestinal organoids (HIOs) derived from human pluripotent stem cells co-differentiate both epithelial and mesenchymal lineages *in vitro* but lack important cell types such as neurons, endothelial cells, and smooth muscle. Here, we report an *in vitro* method to derive HIOs with epithelium, mesenchyme, enteric neuroglial populations, endothelial cells, and organized smooth muscle in a single differentiation, without the need for co-culture. When transplanted into a murine host, these populations expand and organize to support organoid maturation and function. Functional experiments demonstrate enteric nervous system function, with HIOs undergoing peristaltic-like contractions, suggesting the development of a functional neuromuscular unit. HIOs also form functional vasculature, demonstrated *in vitro* using microfluidic devices to introduce vascular-like flow, and *in vivo* following transplantation, where HIO endothelial cells anastomose with host vasculature. Collectively, we report an *in vitro* model of the human gut that simultaneously co-differentiates epithelial, stromal, endothelial, neural, and organized muscle populations.

## Main Text

Human intestinal organoids (HIOs) are three-dimensional structures derived from induced pluripotent stem cells (iPSCs) that model the human intestine^1^ and are powerful tools for studying human development, physiology and pathophysiology^2–5^. Unlike patient-derived organoids (also known as enteroids), HIOs possess cells from multiple tissue lineages, including endoderm-derived epithelial cells and mesoderm-derived mesenchymal cells^1,6^. HIOs are typically simple structures *in vitro,* lacking complex organization and important cell types crucial to intestinal function such as neurons, endothelial cells and organized smooth muscle. This simplicity renders the organoids incapable of recapitulating many functions of the intestine^7^. Several approaches have been developed to increase HIO complexity, including adding missing cell types individually or by co-culture, or transplantation into a murine host where they are vascularized by the host and undergo maturation, developing crypts and villi along with the muscularis mucosae and outer muscle layers, representing a miniature human intestine^6,8,9^. Altering growth factors present during differentiation has also been used as a method to induce additional lineages^10–14^; however, a single, harmonized method to achieve coordinated differentiation of multiple cell lineages in the same organoid has not previously been reported.

Here we leverage recent discoveries that the EGF-family member, EPIREGULIN (EREG), is an important stem cell niche factor in the developing human intestine^15^, and demonstrate that EREG promotes iPSC-derived HIOs containing epithelium, stroma, neurons, endothelial cells, and organized smooth muscle, similar to the native human intestine. Once transplanted into a murine host, these cell populations further organize along a crypt-villus axis. We show that these HIOs can recapitulate important functions of the human intestine as they spontaneously and rhythmically contract without stimulus, suggesting functional muscle and neuronal connections, and can be connected to simulated and real circulatory systems, confirming blood vessel function.

## Results

### Generation of human intestinal organoids containing neurons, endothelial cells, and organized smooth muscle

The directed differentiation method to generate HIOs from human iPSCs is robust and has been widely used for over a decade^1,16^. Broadly, this differentiation approach relies on the induction of a mixed endoderm-mesoderm culture followed by further differentiation into intestinal lineages. During intestinal lineage differentiation, cells growing in 2-dimensional (2D) culture self-organize and form 3-dimensional (3D) spheroids that possess cells derived from endoderm and mesoderm. Spheroids are typically placed in media containing EGF, NOGGIN, and RSPONDIN-1 (ENR Media) for 3 days, which patterns a proximal, small-intestinal identity^11,17^, and can then be cultured in media containing only EGF. These organoids have been extensively characterized by our group and others^1,6,11,18^; single cell RNA-sequencing (scRNA-seq) has revealed that small populations of neuron-like and endothelial-like cells are present in early cultures, but these populations are transient^11,19^. Smooth muscle cells can be found in these organoids, but they are rare, sparsely distributed and unorganized within the HIOs *in vitro*^20^. We recently reported that EPIREGULIN (EREG) is a stem cell niche factor *in vivo* during human intestinal development, and can replace EGF in tissue-derived intestinal enteroid culture, leading to improved spatial organization and enhanced differentiation of all epithelial cell types^15^. A more comprehensive analysis of our previously published scRNA-seq data followed by validation with fluorescence *in situ* hybridization (FISH), shows that EREG is also expressed in the outer longitudinal and circular smooth muscle of the developing human intestine at all time points examined (Extended Data Fig. 1a-e). Based on these new observations, and our prior findings that EGF is expressed in the differentiated villus epithelial cells, we hypothesized that EREG may play a role in mesenchymal patterning and differentiation during intestinal development.

To test if EREG plays a role during HIO differentiation, we followed the previously described differentiation paradigm (Fig. 1a) up to the spheroid stage^1,16^. Once spheroids are collected and embedded in Matrigel, they are patterned for 3 days with ENR media, and subsequently cultured in EGF-only media, at a standard concentration of 100 ng/mL. To determine if EREG can replace EGF in culture and retain differentiation potential, we cultured spheroids in three concentrations of EREG (1 ng/mL, 10 ng/mL, 100 ng/mL) and compared this to EGF (1 ng/mL, 10 ng/mL, 100 ng/mL), with a constant concentration of NOGGIN and RSPO1 for the first three days, before switching to EREG-only or EGF-only media (Fig. 1a).

**Figure 1:**
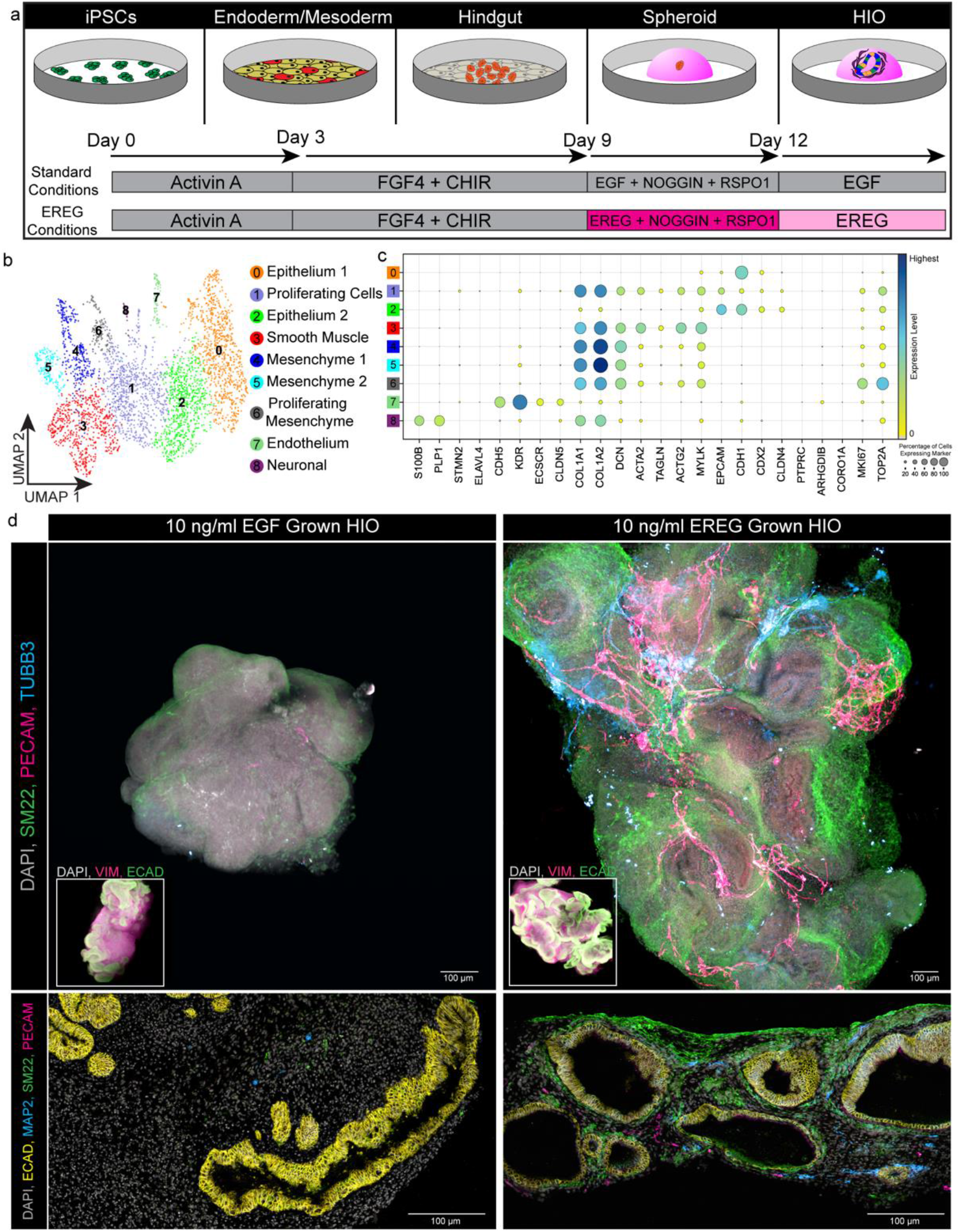
EREG-HIOs grown *in vitro* spontaneously and simultaneously pattern endothelium, smooth muscle, and neural components. a) Schematic of HIO directed differentiation using standard EGF conditions (grey) and experimental EREG conditions (pink). b) UMAP visualization of snRNA-seq from 28-day *in vitro* grown EREG-HIOs in 10 ng/mL of EREG (n=1 sequencing run of over 20 combined HIOs). c) Dot plot visualization for expression of canonical markers of neurons (*S100B, PLP1, STMN2, ELAVL4*), endothelial cells (*CDH5, KDR, ECSCR, CLDN5*), mesenchyme (*COL1A1, COL1A2, DCN*), smooth muscle (*ACTA2, TAGLN, ACTG2, MYLK*), epithelium (*EPCAM, CDH1, CDX2, CLDN4*) immune cells (*PTPRC, ARHGDIB, CORO1A*) and proliferative cells (*MKI67, TOP2A*), in EREG-grown (10 ng/mL) HIOs. d) Top panels: representative whole mount immunofluorescence (IF) staining of 10 ng/mL EGF- or EREG-grown HIOs for the presence of smooth muscle (SM22; green), endothelial cells (PECAM; pink), and neurons (TUBB3; blue). Inlays show IF staining of mesenchyme (VIM; pink) and epithelium (ECAD; green). Bottom panels: representative IF staining on 2D sections of EREG-grown (10 ng/mL) and EGF-grown (10 ng/mlL) HIOs for the presence of epithelium (ECAD; green), smooth muscle (SM22; blue), endothelial cells (PECAM; yellow), and neurons (MAP2; pink). All Scale bars = 100 µm.

We determined the HIO forming efficiency of each concentration by counting the number of spheroids plated in a single well at day 0, and again counting the number of HIOs present after 10 days in culture (formula: # of day 10 HIOs/# of day 0 spheroids=forming efficiency). Experiments were repeated for 3 separate experiments (batches), in 3 different iPSC lines with 3 technical replicates. We saw minimal difference in HIO forming efficiency when comparing ligand or concentration, with growth in 10 ng/mL EREG having the only statistically significant increase in forming efficiency across conditions compared to the EGF control cultures (Extended Data Fig. 2a). We compared morphologic features between the culture conditions, including area, circularity, roundness, solidity, and aspect ratio. In general, we did not see any consistent patterns or major differences in these measurements between doses or media (Extended Data Fig. 2b-f), suggesting all EREG concentrations form HIOs similar in shape and size as the EGF controls.

To characterize HIOs, we subjected all three concentrations of EREG-grown HIOs to scRNA-seq (Extended Data Fig. 3a-b) and found a cluster enriched with smooth muscle genes (cluster 2: *ACTA2*, *TAGLN*, *ACTG2*, *MYLK*) and 3 clusters with neuroglial identities (clusters 5, 7, and 8: *S100B, PLP1*, *STMN2*, *ELAVL4*); which have been found in rare, transient populations in EGF-cultured HIOs previously^11,12,18^ (Extended Data Fig. 3c). When we plotted cluster abundance per sample, we found that most of the cells contributing to smooth muscle and neural clusters were from the EREG samples, with the 1 ng/mL and 10 ng/mL samples having the highest contributions to these lineages (Extended Data Fig. 3d). The 10 ng/mL EREG condition possessed both glial and neuronal populations (Extended Data Fig. 3e-f) and featured the most heterogeneous enteric neuron populations (Extended Data Fig. 3g). Based on the high HIO forming efficiency (Extended Data Fig. 2a) and the robust neural cell types found within the 10 ng/mL EREG condition, we chose this condition for further characterization.

We confirmed the enhanced differentiation of neurons and smooth muscle in EREG culture compared to EGF culture, across three different experiments, and in three different iPSC lines using RT-qPCR (Extended Data Fig. 3h). During this validation, we also observed endothelial gene expression signatures (*CDH5, VEGF*) in all 3 biological replicates examined by RT-qPCR (Extended Data Fig. 3h). Interestingly, no endothelial cells were observed in the whole cell scRNA-seq data. However, endothelial cells are known to be sensitive to dissociation methods and are often lost during preparation of samples for whole cell sequencing^21^. Thus, we isolated nuclei from the 10 ng/mL EREG HIOs and generated single nuclear RNA sequencing (snRNA-seq) data, which supported the presence of endothelial cells (Fig. 1b-c). Notably, fewer neurons were present in snRNA-seq data, as previously reported^22,23^. We next validated the presence of neural, endothelial, and smooth muscle lineages with whole mount 3D immunostaining and in 2D sections of HIOs grown in EGF or EREG, and observed that EREG-grown, but not EGF-grown HIOs, possessed organized SM22^+^ (TAGLN^+^) smooth muscle, networks of TUBB3^+^ (3D) and MAP2^+^ (2D) neuron-like cells, and PECAM^+^ endothelial cells throughout the HIOs (Fig. 1d).

### Transplantation of EREG-HIOs leads to increased maturation and cellular organization

Murine kidney capsule engraftment is well established as a system to further mature *in vitro* grown HIOs^2,5,13,20,24,25^. In short, HIOs grown *in vitro* for 28 days or more are surgically placed under the kidney capsule of immunocompromised NSG mice, where they engraft and grow for an additional 10-12 weeks. These organoids greatly expand in size, mature to become spatially organized similar to the native human intestine, have increased cell type diversity, and can be harvested for snRNA-sequencing and staining^19^ (Fig. 2a). To determine if EREG-grown HIOs could be further matured and become functional, we transplanted 10 ng/mL EREG- or 10 ng/mL EGF-grown control HIOs beneath the kidney capsule and allowed them to grow for 12 weeks. Immunofluorescent (IF) staining for epithelial (ECAD) and smooth muscle (TAGLN) markers revealed that both EGF-transplanted HIOs (tHIOs) and EREG-tHIOs were spatially organized as previously reported with the development of crypt-villus units and organized smooth muscle (Fig. 2b). We observed very few neurons or endothelial cells in control EGF-tHIOs (Fig. 2b, middle panels), while EREG-tHIOs possessed abundant TUBB3^+^ neurons and PECAM^+^ endothelial cells spatially organized similar to the native human intestine (Fig. 2b).

**Figure 2:**
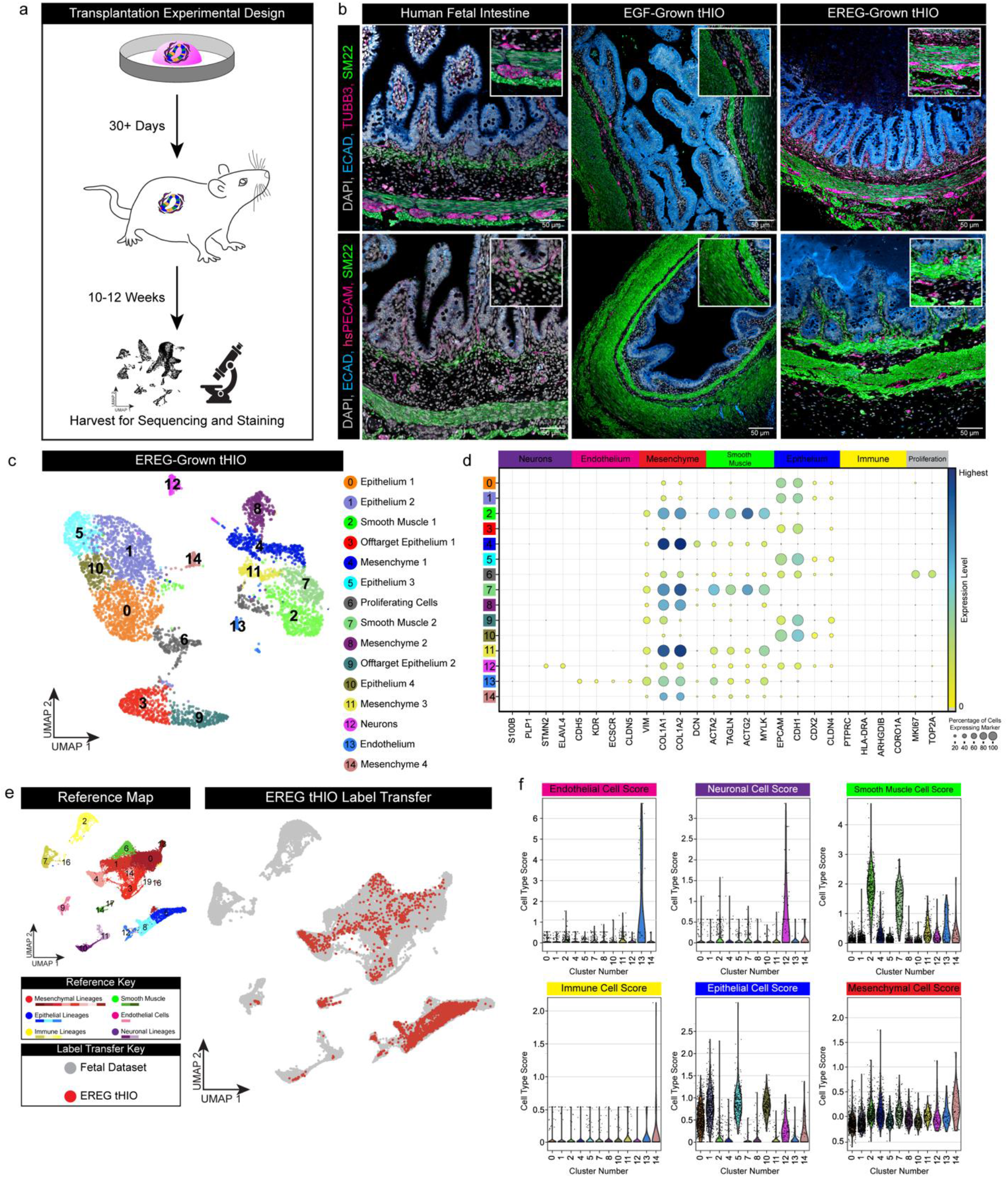
EREG-grown HIOs further mature and spatially organize after transplantation into murine kidney capsule. a) Schematic timeline of HIO transplantation experiment. b) Representative IF staining of human fetal intestine (left panel; 127 days post conception), EGF-grown tHIO (middle panel; 12 weeks), and EREG-grown tHIO (right panel; 12 weeks) stained for the presence of smooth muscle (SM22; green), epithelium (ECAD; blue), and neurons (TUBB3; pink) in top panels. Bottom panels show stains for the presence of smooth muscle (SM22; green), epithelium (ECAD; blue), and endothelial cells (PECAM; pink). All scale bars = 50 µm. c) UMAP visualization of snRNA-seq of 12-week *in vivo* grown tHIOs in 10ng/mL of EREG (n=1 sequencing run of one tHIO). d) Dot plot visualization for expression of canonical markers of neurons (*S100B, PLP1, STMN2, ELAVL4*), endothelial cells (*CDH5, KDR, ECSCR, CLDN5*), mesenchyme (VIM, COL1A1, COL1A2, DCN), smooth muscle (*ACTA2, TAGLN, ACTG2, MYLK*), epithelium (*EPCAM, CDH1, CDX2, CLDN4*) immune cells (*PTPRC, HLA-DRA, ARHGDIB, CORO1A*) and proliferative cells (*MKI67, TOP2A*). e) Left: UMAP visualization of human fetal intestinal data set from Extended Data Fig. 1 recolored for cell type lineages: mesenchymal cells(red), epithelial cells (blue), immune cells (yellow), smooth muscle cells (green), endothelial cells (pink), and neuronal cells (purple). Right: UMAP visualization of label transfer results with reference human fetal intestinal dataset in grey and tHIO dataset in red. f) Violin plot quantification of cell type scoring for each tissue type in reference intestinal dataset.

SnRNA-sequencing of EREG-tHIOs confirmed the presence of epithelial cells, mesenchymal cells/fibroblasts, smooth muscle, neural cells, and endothelial cells (Fig. 2c-d). We found two undefined epithelial clusters (clusters 3 and 9) appeared to be mixed gastric and intestinal epithelium based on mapping to a human fetal endoderm atlas^19^ (Extended Data Fig. 4a). Small portions of undefined or non-intestinal cells have previously been reported in HIOs^19^, and these cells were excluded from further analysis. To confirm cluster annotations based on marker gene expression (Fig. 2d), we carried out a label transfer of EREG-tHIO data onto a previously published fetal intestine reference dataset^26^ (Fig. 2e and Extended Data Fig. 1a), and confirmed the accuracy of annotations based on marker genes. tHIO datasets robustly label transferred onto their counterparts in the human fetal datasets using Seurat’s integrated label transfer function^27^ (Fig. 2e, Extended Data Fig. 4b). To further benchmark tHIO cell type identity against our reference dataset, we used a cohort of highly enriched genes for each major cell type in the reference (endothelial, neural, smooth muscle, immune, epithelial, and mesenchymal), that allowed us to generate a “score” for each cell in the tHIO sample based on the enrichment of these genes. Cluster 6 contained both proliferating mesenchymal and epithelial cells, so we excluded it from this analysis for clarity. We found that the annotated tHIO clusters scored highly for their counterparts in the reference set, further confirming our annotations and the label transfer findings (Fig. 2f, Extended Data Fig. 4b, Extended Data Table 1). Collectively, this data suggests the cell types within tHIOs, including endothelial cells and neurons, share transcriptional states with their analogous cell type within the human intestine.

### EREG-tHIOs exhibit peristaltic-like function

To test if the neurons and smooth muscle seen within EREG-tHIOs create a functional neuromuscular unit, we dissected control EGF-tHIOs and EREG-tHIOs into small muscle strips and explanted them into Kreb’s buffer in an organ bath chamber to measure muscle contractile force, as previously reported^12,28^. Explanted muscle strips were allowed to equilibrate and monitored continuously for contractions (Fig. 3a). In the absence of any stimulation, we observed spontaneous and rhythmic contractions in EREG-tHIOs, a phenomenon that was not seen in control EGF-tHIOs (Fig. 3b). These phasic contractions suggest the presence of intramuscular interstitial cells of Cajal (ICCs) in EREG-grown tHIOs^12,29^. We stained for the neuronal marker TUBB3 and the ICC marker c-KIT and found that EGF-tHIOs had rare TUBB3^+^ neurons and completely lacked c-KIT^+^ ICCs. On the other hand, EREG-tHIOs contained many c-KIT^+^ ICCs directly adjacent to TUBB3^+^ neurons, closely resembling the staining pattern found in the neural plexus of the developing human intestine (Fig. 3c).

**Figure 3:**
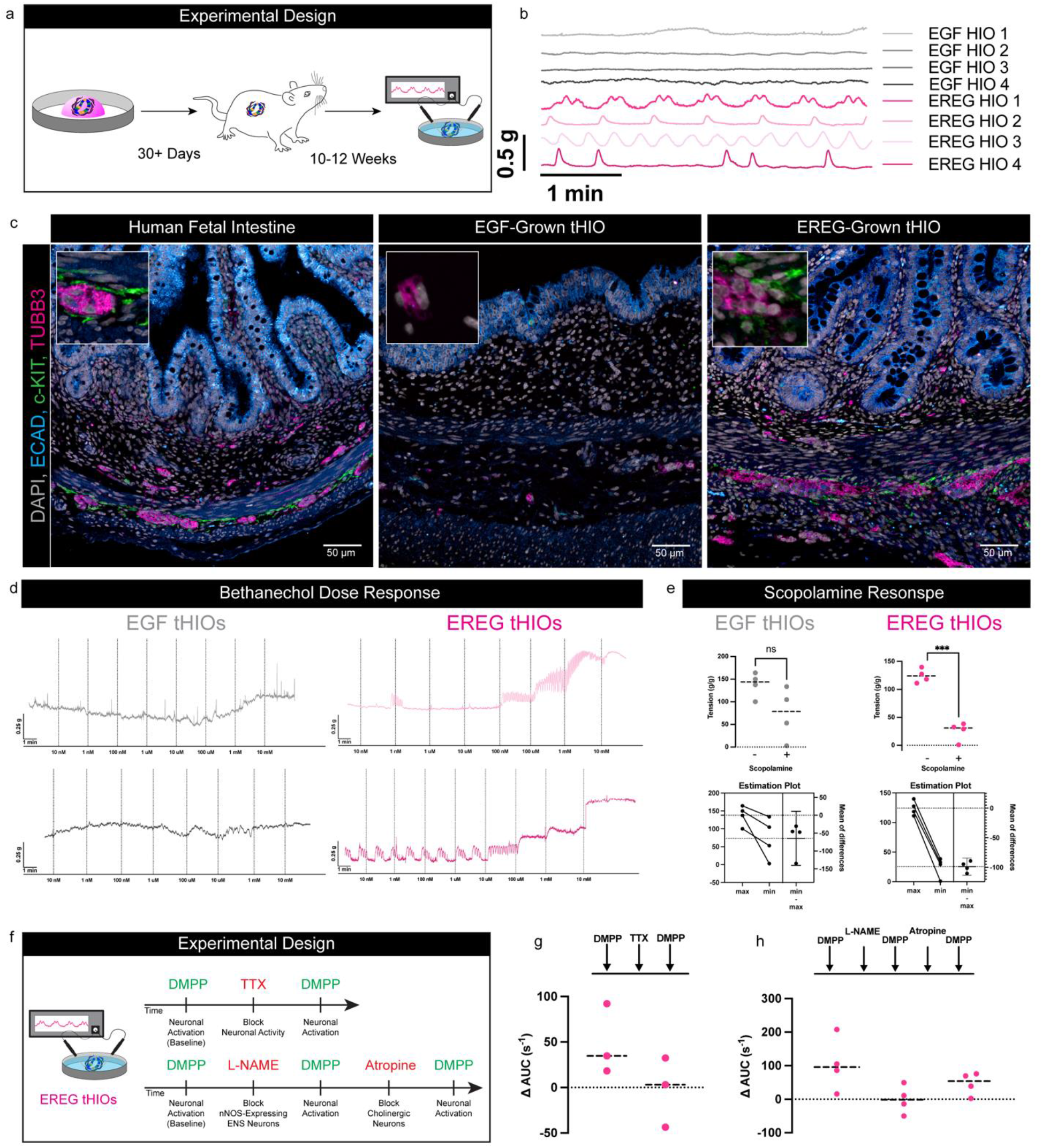
Assessment of EREG-grown tHIOs for neuromuscular units and native functionality. a) Experimental schematic for transplanting *in vitro* grown HIOs under the kidney capsule of a murine host and testing muscular and ENS function in an organ bath measured with isometric-force transducers post-transplant. b) Isometric force contractions in tissues isolated from n=4 different EGF-grown tHIOs (grey) and four different EREG-grown tHIOs (pink) after an equilibrium period with no exogenous contractile triggers. c) Representative IF staining of human fetal intestine (Left; 127 days post conception), EGF-grown tHIO (Middle; 12-weeks), EREG-grown tHIO (Right; 12-weeks) stained for the presence of epithelium (ECAD; blue), general neurons (TUBB3; pink) and ICC’s (c-KIT; green). All scale bars = 50 µm. d) Activation of muscarinic receptor-induced contractions in tissues isolated from n=2 EGF-grown tHIOs (grey) and n=2 EREG-grown tHIOs (pink) using increasing doses of bethanechol. e) Inhibition of the muscarinic receptor with scopolamine induced muscle relaxation. Graphs show calculated maximum and minimum tissue tension from n=2 EGF-grown tHIOs (grey) and n=2 EREG-grown tHIOs (pink). f) Functional test of ENS inhibition using the neurotoxin tetrodotoxin (TTX). Addition of TTX lowers ENS activation in the presence of DMPP stimulation. Graphed is the change in AUC following a control DMPP stimulation measured after stimulation, followed by TTX treatment and a final DMPP stimulation in EREG-grown tHIOs. g) Functional test of specific ENS neuronal types (nitrergic and cholinergic) in muscle contractions. Inhibition using the nitrergic inhibitor L-NAME and the cholinergic inhibitor atropine. Graphed is the change in AUC following a control DMPP stimulation measured after stimulation, followed by L-NAME treatment, another DMPP stimulation, followed by Atropine treatment, and a final DMPP stimulation in EREG-grown tHIOs.

Next, we treated the organoids with bethanechol, a muscarinic receptor agonist that directly stimulates muscle contractions, and measured contractile force. No notable change in contractile force could be measured in control EGF-tHIOs (Fig. 3d – left, grey) while contractile force increased in a dose-dependent manner in response to bethanechol in EREG-tHIOs (Fig. 3d – right, pink). We then treated the explanted tHIOs with scopolamine, a muscarinic antagonist that blocks smooth muscle contraction, and were able to trigger significant muscle relaxation (Fig. 3e - right, pink) in EREG-tHIOs but not in EGF-tHIOs (Fig. 3e – left, grey). With these data suggesting functional muscle and the presence of neurons in these organoids, we hypothesized the presence of a functional neuromuscular unit in EREG-tHIOs.

To examine if the neuronal populations functionally regulate smooth muscle contractions, we excited neurons by using the selective α3-nicotinic receptor agonist dimethylphenylpiperazinium (DMPP) to stimulate neurotransmitter release and activate neurons in EREG-grown tHIOs (Fig. 3f). Following DMPP treatment, explants were treated with tetrodotoxin (TTX) to block action potentials which successfully blocked the neurons’ ability to be depolarized again following another dose of DMPP, thus supporting ENS-dependent contractile activity within the tissue (Fig. 3g). Finally, we assessed the function of nNOS-expressing neurons by inhibiting them with NG-nitro-L-arginine methyl ester (L-NAME), and cholinergic neurons by blocking them with a dose of atropine. We then measured contractile activity following a baseline stimulation, or stimulation after exposure to either inhibitor (Fig. 3h). Contractile activity was measured as the change in the area under the curve (ΔAUC) immediately before and after each stimulation. After each inhibitor was added, we saw a significant decrease in muscle relaxation compared to the uninhibited contractions (Fig. 3h), suggesting nNOS-expressing neurons and cholinergic neurons elicit smooth contractions in EREG-tHIO explants. These data together demonstrate EREG-tHIOs not only possess glial, neuronal, and smooth muscle populations but that these populations are functional and collectively drive peristaltic smooth muscle-like contractions.

### EREG-HIOs possess endothelial cells that organize into functional vasculature

Tissue-specific endothelial cells are a critical cell type in all organs as they not only support the metabolic demands of a particular organ, but also supply paracrine angiocrine factors that orchestrate organ development, repair, and regeneration. Corroborating our single nucleus data (Fig. 1b-c), we used whole mount IF staining to interrogate many individual EREG-HIOs and consistently observed robust endothelial cells networks throughout the organoids (Fig. 4a-b). As with the presence of endogenous neurons and smooth muscle structures, we were interested in understanding if these endothelial cells could form functional vessels.

**Figure 4:**
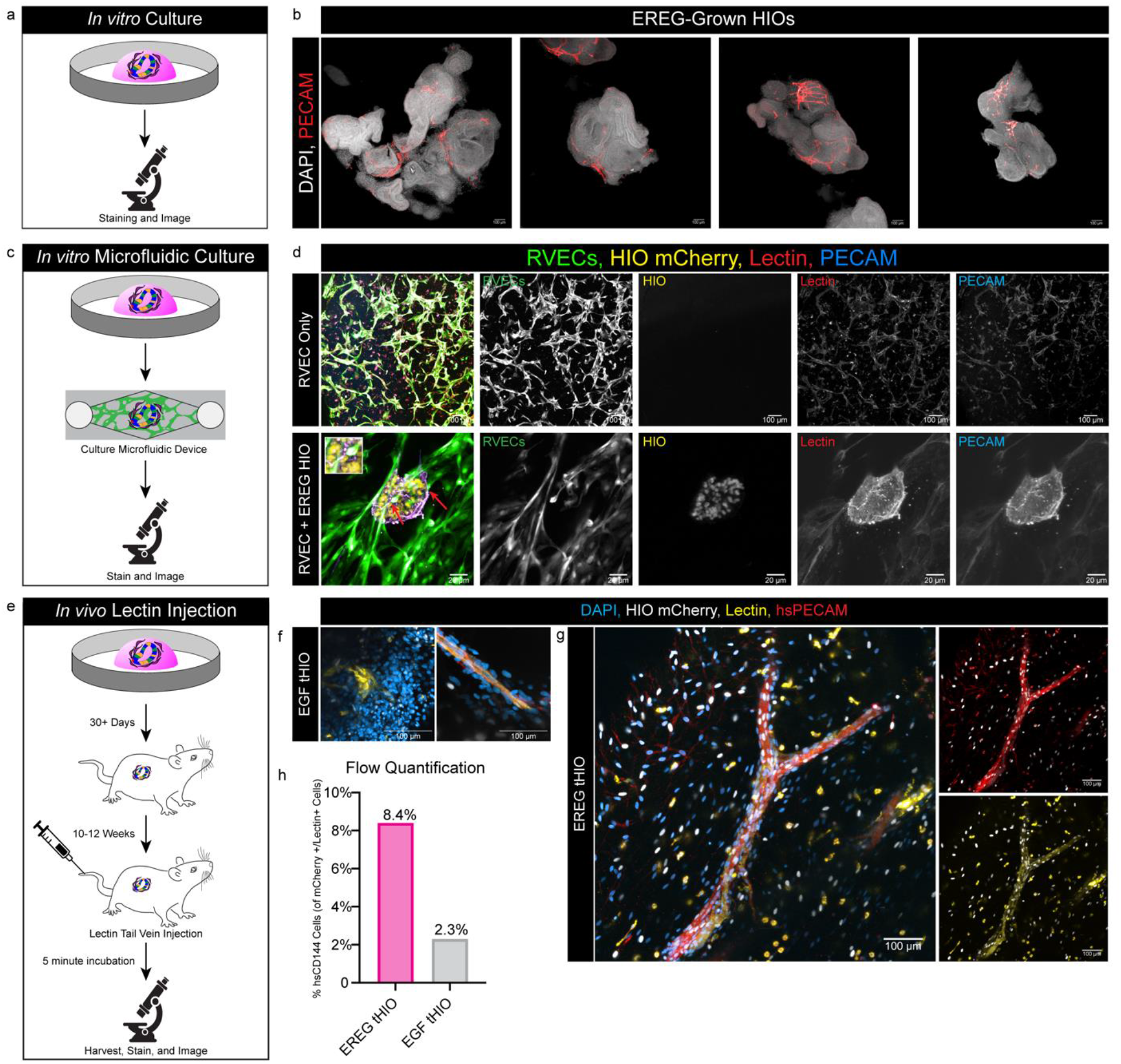
EREG-grown HIOs pattern blood vessels that are functional both *in vitro* and *in vivo.* a) Schematic of workflow for whole mount imaging *in vitro* EREG*-*grown HIOs. b) Representative IF staining of n=4 different EREG-grown (10 ng/mL) HIOs for the presence of endothelial cells (PECAM; red), and DAPI (grey). All scale bars = 100 µm. c) Schematic of workflow for *in vitro* EREG*-*grown HIO functionality test using RVEC microfluidic device. d) Representative IF staining of the control RVEC only microfluidic device lane (top) and the RVEC + EREG-grown (10 ng/mL) HIOs lane. RVECs (green), mCherry^+^ HIOs (yellow), lectin dye flown through system (red) and PECAM dye flown through system (blue). Overlap (purple) in HIOs are areas where RVECs connected with endogenous endothelial cells and lectin flow moved through vessels. Control scale bars = 100 µm and RVEC + HIO image scale bars = 20 µm. e) Schematic of workflow for *in vivo* EREG*-*grown tHIO functionality test for connection with host vasculature. f) Representative whole mount IF staining of EGF-grown (10 ng/mL) tHIOs for the presence of human endothelial cells (hsPECAM; red), HIO mCherry tag (white), lectin dye administered through tail vein injection (yellow) and DAPI (blue). All scale bars = 100 µm. g) Representative whole mount IF staining of EREG-grown (10 ng/mL) tHIOs for the presence of human endothelial cells (hsPECAM; red), HIO mCherry tag (white), lectin dye administered through tail vein injection (yellow) and DAPI (blue). All scale bars = 100 µm. h) Quantification of flow cytometry analysis to quantify the percentage of hsPECAM^+^/Lectin^+^ cells. Three 12-week-old tHIOs per condition were pooled per condition to ensure enough material for experiment.

To test their tubulogenic capabilities *in vitro,* we leveraged human ‘reset-vascular endothelial cells’ (RVECs) which have been shown to self-assemble into stable, multilayered and branching perfusable, vascular networks within scalable microfluidic chambers, which are capable of transporting human blood, and vascularizing colonic enteroids^30^ (Fig. 4c). R-VECs are engineered by transient introduction of the pioneer transcription factor ETV2 into human endothelial cells, conferring them with the capacity to respond to biophysical and biochemical signals emanating from the microenvironment, such as intestinal epithelial cells. We hypothesized that RVECs would adapt and anastomose to the endogenous endothelial cells within the EREG-HIO, enabling flow through the HIO. To this end, iPSCs labeled with a lentivirus expressing nuclear mCherry were used to derive mCherry^+^ EREG-HIOs. After 14 days in culture, mCherry^+^ EREG-HIOs were mixed with GFP-labeled RVECs in a fibrin matrix and seeded into a microfluidic device. After 48 hours, a tomato lectin dye and PECAM antibody were flowed through the system to label any vascular networks that had formed^31^. We observed GFP^+^ RVECs anastomosing to mCherry^+^/PECAM^+^/Lectin^+^ networks within the EREG-HIOs suggesting that endogenous endothelial cells were not only able to connect to the RVEC network but that they formed perfusable vessel-like structures that enabled flow of media and lectin (Fig. 4d).

To interrogate the function of endothelial cells within EREG-HIOs *in vivo*, we transplanted EREG-HIOs under the kidney capsule of a mouse and allowed them to grow for 10 weeks. These organoids were harvested and co-stained with a human-specific PECAM (hsPECAM) antibody and a pan-VE-CAD antibody that cross reacts with both human and mouse to confirm species specificity of observed endothelial cells (Extended Data Fig. 5a). Within EGF-tHIOs, we noticed only rare human endothelial cells labeled with the hsPECAM antibody. In contrast, EREG-tHIOs contained abundant hsPECAM labeling (Extended Data Fig. 5b). We also observed that murine red blood cells autofluoresce in the 488-channel, so we leveraged this autofluorescence to show that hsPECAM^+^ vascular structures within EREG-tHIOs were filled with mouse red blood cells (Extended Data Fig. 5c), strongly suggesting that EREG-tHIO endothelial cells anastomose with the host’s circulatory system. Based on hsPECAM/VE-CAD co-labeling, we observed multiple points where a mouse blood vessel (VE-CAD^+^/hsPECAM^neg^), appeared to connect to a human blood vessel (PECAM^+^/VE-CAD^+^) further suggesting that EREG-tHIO endothelial cells anastomose with the mouse circulatory system to form functional vessels (Extended Data Fig. 5d).

To directly test a functional connection between the murine and EREG-tHIO vasculature, we transplanted organoids into a murine host and allowed them to mature for 10 weeks (Fig. 4e). We then injected the host with tomato-lectin via tail vein injection^30^ and allowed the lectin to circulate for 5 minutes. Organoids were then harvested and whole mount IF stained with the hsPECAM antibody to delineate between mouse and human endothelial cells (Fig. 4e). Within EGF-tHIOs, we were only able to find small rare Lectin^+^ staining of vascular-like structures, which were largely hsPECAM-negative, suggesting that most vessels in these tHIOs were host-derived (Fig. 4f). On the other hand, in EREG-tHIOs, large mCherry^+^/hsPECAM^+^/Lectin^+^ vessels were readily observed (Fig. 4g). Lectin staining in mCherry^+^/hsPECAM^+^ structures suggest these human vessels are connected to the host’s circulatory system. To quantify the proportion of human-specific endothelial cells labeled by lectin, we dissociated these transplanted organoids after lectin injection and used flow cytometry to determine the proportion of mCherry^+^ human cells that were also positive for a VE-CAD flow antibody (CD144)^+^/Lectin^+^ in both EGF-tHIOs and EREG-tHIOs. We found that EREG-tHIOs had a much larger proportion of human blood vessels labeled with lectin (8.4%) compared to EGF-tHIOs (2.3%) (Fig. 4h and Extended Data Fig. 6a-b). These data taken together demonstrate EREG-grown HIO’s endogenous endothelial population is able to connect to circulatory systems both *in vitro* and *in vivo* and have the ability to enable function perfusion to sustain their viability and function.

## Discussion

By leveraging information from the developing human intestine, which allowed us to identify the developmental niche factor EREG, we have generated the first hPSC-derived human intestinal organoid model that can simultaneously pattern epithelial, mesenchymal/stromal, neural, endothelial and smooth muscle populations. By transplanting these organoids into a murine host, we were able to expand and mature HIOs further in order to assess function. EREG-tHIOs not only contain neurons and endothelial cell populations, but they also function *in vivo.* Functional experiments to interrogate smooth muscle contractions strongly support the development of a functional neuromuscular unit between neural populations and smooth muscle populations; similarly, several lines of evidence support that HIO-derived endothelial cells within the organoid connect with *in vitro* and *in vivo* circulatory systems, creating vasculature capable of flow. While our findings are promising, enhancing HIO differentiation with previously missing lineages, making significant progress towards creating a fully artificial intestine *in vitro,* we note that there is still room for progress and improvement, given that there are still missing cell types such as immune and lymphatic lineages, which play critical roles in the intestinal function and disease.

Taken together, our results demonstrate the ability to create more physiologically relevant culture conditions to pattern complex and accurate organoid models of the human intestine. This system improves complexity and may enhance the utility of HIOs to understand human intestinal development, disease modeling, drug screening, and personalized medicine.

## Methods

### Microscopy

All fluorescence images were taken on a Nikon AXR confocal microscope. Acquisition parameters were kept consistent within the same experiment and all post-image processing was performed equally across all images of the same experiment. Images were assembled in Adobe Photoshop CC 2023, and Figures were assembled using Adobe Illustrator CC 2023.

### Tissue Processing and Staining

#### Tissue processing for staining and histology

All tissue or HIOs were placed in 10% Neutral Buffered Formalin (NBF) for 24 hours at room temperature (RT) on a rocker for fixation. Fixed specimens were then washed 3x in UltraPure DNase/RNase-Free Distilled Water (Thermo Fisher Cat #10977015) for 30-60 minutes per wash depending on its size. Next, tissue was dehydrated through a methanol series diluted in UltraPure DNase/RNase-Free Distilled Water for 30-60 minutes per solution: 25% MeOH, 50% MeOH, 75% MeOH, 100% MeOH. Tissue was either immediately processed for paraffin embedding or stored in 100% MeOH at 4°C for future paraffin processing or whole mount staining. For paraffin processing, dehydrated tissue was placed in 100% EtOH, followed by 70% EtOH, and perfused with paraffin using an automated tissue processor (Leica ASP300) with 1 hour solution changes overnight. Tissue was then placed into tissue cassettes and base molds for sectioning. Prior to sectioning, the microtome and slides were sprayed with RNase Away (Thermo Fisher Cat#700511). 5 µm-thick sections were cut from paraffin blocks onto charged glass slides. Slides were baked for 1 hour at 60°C in a dry oven and were used within 24 hours for FISH or within a week for IF. Slides were stored at room temperature in a slide box containing a silica desiccator packet and the slide box seams were sealed with parafilm.

#### Antibody, Fluorescence in situ hybridization (FISH) Probes, and dye information

All stains were completed in organoids derived from 3 different stem cell lines, representative images from one line (iPSC72.3) are reported in this manuscript. The following antibodies were used throughout the manuscript in immunofluorescence and fluorescent *in situ* hybridization with co-immunofluorescence staining. All antibodies were used on FFPE processed sections or whole mount organoids as described below, no frozen sections were used. Rabbit anti-SM22 1:500 (Abcam Cat#ab14106), goat anti-E-Cadherin 1:500 (R&D Systems Cat# AF748), mouse anti-E-Cadherin 1:500 (BD Transduction Laboratories Cat# 610181), rabbit anti-PECAM 1:200 (Sigma Cat# HPA004690), mouse anti-TUBB3 1:200 (BioLegend Cat# 801201), sheep anti-SM22 1:500 (Novus Cat# AF7886), chicken anti-MAP2 1:2000 (Abcam Cat#ab92434), goat anti-VIM 1:200 (R&D Systems Cat#AF2105), rabbit anti-c-KIT 1:500 (Abcam Cat#ab32363) and mouse anti-VE-CAD 1:1000 (R&D Systems Cat#MAB9381). FISH probes were acquired from ACDbio and stained using the RNAscope multiplex fluorescent manual protocol and kit.

RNAscope Probe Hs-EREG (ACD, Cat# 313081). Tomato lectin dye was purchased from Thermo Fischer Cat#L32472 and was used 1:1 with sterile PBS for tail vein injections and diluted 1:100 in Tube media (See “RVEC Experiments” section below for Tube Media formulation) for RVEC experiments.

#### Immunofluorescence (IF) protein staining on 2D paraffin sections

Tissue slides were deparaffinized in Histo-Clear II (National Diagnostics Cat#HS-202) twice for 5 minutes each, followed by rehydration through an ethanol series of two washes each for two minutes in 100% EtOH, 95% EtOH, 70% EtOH, 30% EtOH, and finally two washes in ddH2O each for 5 minutes. Antigen retrieval was performed using 1X Sodium Citrate Buffer (100mM trisodium citrate (Sigma, Cat#S1804), and 0.5% Tween 20 (Thermo Fisher Cat#BP337), pH 6.0). Slides were steamed for 20 minutes then washed three times in ddH2O for 5 minutes each. Slides were then incubated in a humidity chamber at room temperature for 1 hour with blocking solution covering the tissue (5% normal donkey serum (Sigma Cat#D9663) in PBS with 0.1% Tween 20). Slides were then incubated in primary antibody diluted as stated above in blocking solution at 4°C overnight in a humidity chamber. The next day, slides were washed three times in 1X PBS for 5 minutes each and incubated with secondary antibody (1:500) with DAPI (1:1000) diluted in blocking solution for 1 hour at room temperature in a humidity chamber. Secondary antibodies were raised in donkey and purchased from Jackson Immuno. Slides were then washed 3x in 1X PBS for 5 minutes each and mounted with ProLong Gold (Thermo Fisher Cat#P369300). Immunofluorescent stains were imaged within 2 weeks. Stained slides were stored flat and in the dark at 4°C.

#### FISH on 2D paraffin sections

FISH staining protocol was performed according to the manufacturer’s instructions (ACDbio, RNAscope multiplex fluorescent manual) with a 30 minute protease treatment and a 20 minute antigen retrieval step. IF protein co-stains were added following the ACDBio FISH protocol. Briefly, after blocker is applied to the final channel and washed twice in wash buffer, slides were washed 3x for 5 minutes in PBS followed by the IF protocol stated above from the blocking step onwards. FISH stains were imaged within a week.

#### Whole mount IF with antibody staining

Organoids were removed from Matrigel using a cut P1000 tip and transferred to a 1.5 mL mini-centrifuge tube. Tubes were spun at 300g for 5 minutes at 4°C and supernatant was removed. Organoids were fixed in 10% NBF overnight at room temperature on a rocker. The following day, organoids were washed 3x for 1 hour in organoid wash buffer (OWB) (0.1% Triton, 0.2% BSA in 1× PBS) at room temperature on a rocker. Organoids were then incubated in CUBIC-L (TCI Chemicals Cat#T3740) for 24 hours at 37°C. They were then washed 3x in OWB and permeabilized for 24 hours at 4°C on a rocker with permeabilization solution (5% normal donkey serum, 0.5% Triton in 1× PBS). After 24 hours of permeabilization, the solution was removed and the desired primary antibody, diluted in OWB, was added. Organoids were incubated overnight at 4°C on a rocker. The next day, organoids were washed 3x in OWB for 1 hour per wash at room temperature on a shaker. Then, secondary antibody was diluted in OWB at 1:500 and added overnight at 4°C wrapped in foil on a shaker. Organoids were then washed again the following day 3x in OWB at room temperature with the first wash being for 1 hour with DAPI added at dilution of 1:1,000. The remaining two washes were for 1 hour in OWB only. Organoids were then transferred to a 96-well imaging plate (Thermo Fisher Cat#12-566-70) and cleared using enough CUBIC-R to submerge the organoids (TCI Chemicals Cat#T3741). Organoids remained in CUBIC-R for imaging and whole mount images were imaged within 1 week.

### HIO Cultures

#### Stem cell lines and generation of human intestinal organoids (HIOs)

This study includes data from HIOs generated across 3 hPSC lines: Human ES line H9 (NIH registry #0062, RRID: CVCL_9773, female) with an mCherry reporter, human iPSC lines WTC11 (RRID: CVCL_Y803, male) and 72.3^31^. All experiments using hPSCs were approved by the University of Michigan Human Pluripotent Stem Cell Research Oversight Committee. All stem cell and organoid lines were routinely monitored for mycoplasma using the MycoAlert Mycoplasma Detection Kit (Lonza Cat#LT07-318).

#### Stem cell maintenance and differentiation

Maintenance and differentiation into HIOs were carried out as previously described^1,2,11,16,32,33^. Cells were kept in a 37°C tissue culture incubator with 5% CO2 and lines were maintained in mTeSR Plus cultured media (Stemcell Technologies Cat# 100-1130). Stem cells underwent directed differentiation into definitive endoderm over a 3-day treatment using Activin A (100ng/mL, R&D Systems Cat#338-AC) added to RPMI base media. This base media was supplemented with 0%, 0.2%, 2% HyClone dFBS (Thermo Fischer Cat#SH3007103) on subsequent days with the addition of 5 mL penicillin-streptomycin each day (Gibco Cat# 15070063). After three days, endoderm monolayers were differentiated into an intestinal identity by treatment with FGF4^34^ (500ng/mL) and CHIR99021 (2μM, APExBIO Cat#A8396). On days 4-6 of hindgut differentiation, spheroids budded from the monolayer and were collected. These spheroids were embedded in Matrigel as previously described^32^ and maintained in basal growth media consisting of Advanced DMEM/F12 (Gibco Cat# 11320033) with B27 (50x, Thermo Fisher Cat#17504044), GlutaMAX (1X, Gibco Cat#35050061), penicillin-streptomycin (Gibco Cat# 15070063), and HEPES buffer (15 mM, Gibco Cat#15630080). Organoid basal growth media was supplemented with epidermal growth factor (EGF) (100 ng/mL, 10 ng/mL, 1 ng/mL R&D Systems Cat#236-EG-01M) or Epiregulin (EREG) (100 ng/mL, 10 ng/mL, 1 ng/mL R&D Systems Cat#1195-EP-025/CF) with Noggin-Fc (100ng/mL, purified from conditioned media^35^), and R-Spondin1 (5% conditioned medium^36^) for the first three days of culture to pattern a proximal small intestine. On the third day after embedding, media was changed to basal growth media supplemented with EGF or EREG only (no additional Noggin or R-Spondin1) and remained in this media for the duration of the experiments with media changes every 5 days. Organoids were not passaged to avoid disrupting the development and spatial organization of the key cell types seen in EREG-grown HIOs.

#### HIO forming efficiency assay

Spheroids were collected from three different stem cell lines for three different batches on days 4-6 of hindgut treatment. Spheroids were plated in Matrigel, counted (day 0), and allowed to grow for 10 days into organoids. After 10 days, organoids were counted and forming efficiency was calculated by taking the number of organoids that had formed at day 10 and dividing it by the total number of spheroids collected on day 0.

#### HIO shape and area quantification

To compare shape and area of different HIO conditions, 5 organoids per condition were grown for 30 days *in vitro* and a 10x bright-field image of each organoid was outlined manually using the freehand selection tool in ImageJ. Outlines were measured in ImageJ with measurements set to capture area and shape descriptors including area, solidity, aspect ratio, circularity, and roundness. This was completed on three stem cell lines and measurements were graphed in Extended Data Fig. 2.

#### RNA extraction, cDNA synthesis, and RT-qPCR

Three different stem cell lines were used for each experiment with three different organoid differentiations (batches) and three technical replicates for each batch. mRNA was isolated using the MagMAX-96 Total RNA Isolation Kit/machine (Thermo Fisher Cat#AM1830), and RNA quality/yield were then measured using a NanoDrop™ One^C^ spectrophotometer (Thermo Fisher Cat#13-400-519) prior to cDNA synthesis. cDNA synthesis was performed using 100 ng of RNA from each sample leveraging the SuperScript VILO cDNA Kit (Thermo Fisher Cat#11754250). RT-qPCR was performed on a Step One Plus Real-Time PCR System (Thermo Fisher Cat#43765592R) with QuantiTect SYBR Green PCR Kit (QIAGEN Cat#204145). Expression of genes in the measurement of arbitrary units was calculated relative to RN18S using the following equation and reported in bar graphs for each gene analyzed: 2^RN18S(CT) − GENE(CT)^ × 1,000.

#### Quantification and statistical analysis (for RT-qPCR etc)

All quantitative experiments were completed in 3 different organoid lines for 3 different batches with 3 technical replicates per batch. All statistical analysis was performed in GraphPad Prism Software. See figure legends for number of replicates used, statistical test performed, and the p-values used to determine the significance for each separate analysis. All t tests were ra”n two-tailed, unpaired with welch’s correction.

#### Mouse kidney capsule transplantation

The University of Michigan and Cincinnati Children’s Hospital Institutional Animal Care and Use Committees approved all animal research. HIOs were cultured *in vitro* for at least 28 days then collected for transplantation. HIOs were implanted under the kidney capsules of immunocompromised NOD-scid IL2Rg-null (NSG) mice^23,25^ (Jackson Laboratory strain no. 0005557). Briefly, mice were anesthetized using 2% isoflurane and a left-flank incision was used to expose the kidney after shaving and sterilization of the area of incision with 3 alternating washes of hibiclens surgical soap and sterile water to prep the area after shaving. Between 1 and 3 HIOs were then surgically implanted beneath mouse kidney capsules using forceps. Prior to closure, an intraperitoneal flush of Zosyn (100 mg kg−1; Pfizer) was administered. Mice were administered a dose of analgesic carprofen during the surgery and an additional dose after 24 hours. All mice were monitored daily for 10 days and then weekly until they were euthanized for retrieval of transplanted HIOs after 10 weeks.

#### tHIO vasculature lectin labeling

HIOs were transplanted into the kidney capsule of a mouse as described above and allowed to mature for 10 weeks. At 10 weeks, conjugated 647 tomato lectin (Thermo Fischer Cat#L32472) was mixed 1:1 with sterile PBS and 100 μL was drawn into a 30-gauge insulin needle. Mice were given a tail vein injection of the diluted lectin and allowed to move about normally for 5 minutes before they were sacrificed for tHIO harvest. tHIOs were immediately placed in 10% NBF overnight at room temperature on a shaker and the whole mount staining protocol outlined in the previous section was started the following day.

#### Flow cytometry

After lectin injection outlined in the previous section, tHIOs were harvested and minced with dissecting scissors. Tissue was then placed into a 15mL conical tube containing 9mL 0.1% (w/v) filter-sterilized Collagenase Type II (Thermo Fisher Cat#17101015) in 1X PBS and 1mL filter-sterilized 2.5 units/mL dispase II (Thermo Fisher Cat#17105041) in 1X PBS per gram of tissue. The tube was incubated at 37°C for 30 minutes with mechanical dissociation every 10 minutes. After incubation, 75 μL DNase I was added and incubated at 37°C for an additional 30 minutes with mechanical dissociation every 10 minutes. Following dissociation, 5 mL of isolation media containing 79% RPMI 1640 (Thermo Fisher Cat#11875093), 20% FBS (Sigma Cat#12103C), and 100 U/mL penicillin-streptomycin (Thermo Fisher Cat#15140122) were added per 10 mL of digestion solution. Cells were filtered through 100 μm and 70 μm filters, pre-coated with isolation media, and centrifuged at 500g for 5 minutes at 4°C. The cells were washed by adding 2 mL of FACS buffer and centrifuged at 500g for 5 minutes at 4°C twice. Cells for all control tubes (unstained, DAPI only, isotype controls, individual antibodies/fluorophores) and experimental cells were placed into a FACS tube for cell sorting (Corning Cat#352063). Cells were stained with primary antibody (CD144) diluted 1:50 in FACS buffer (CD144, VE-Cadherin, anti-human FITC) for 30 minutes at 4°C. Cells were then washed with 5 mL FACS buffer and centrifuging at 500g for 5 minutes at 4°C for two washes. Cells were resuspended in FACS buffer and 0.2 μg/mL DAPI was added. FACS was performed using a BD FACS Discovery S8 Cell Sorter and quantitated using the accompanying software.

### Sequencing Experiments

#### Single cell RNA sequencing dissociation

To dissociate HIOs to single cells, organoids were removed from Matrigel using a cut P1000 tip and placed in a 1.5 mL micro-centrifuge tube. All consumables such as tubes and pipette tips used in this prep were pre-washed with 1% BSA in 1X HBSS to prevent adhesion of cells. Following collection, dissociation enzymes and reagents from the Neural Tissue Dissociation Kit (Miltenyi Cat#130-092-628) were used, and all incubation steps were carried out in a refrigerated centrifuge pre-chilled to 10°C unless otherwise stated. Organoids were treated for 15 minutes at 10°C with Mix 1 followed by an incubation for 10 min increments at 10°C with Mix 2. Frequent agitation by pipetting with a P1000 pipette was implemented until organoids were fully dissociated. Cells were passed through a 70 µm filter coated with 1% BSA in 1X HBSS, centrifuged at 500g for 5 minutes at 10°C and resuspended in 500 mL 1X HBSS (with Mg2+, Ca2+). Cells were centrifuged 500g for 5 minutes at 10°C and washed twice by suspension in 2 mL of HBSS + 1% BSA, followed by more centrifugation. Cells were then counted using a hemocytometer, centrifuged and resuspended to reach a concentration of 1000 cells/µL and kept on ice. Single cell libraries were immediately prepared on the 10x Chromium by the University of Michigan Advanced Genomics Core facility with a target capture of 5000 cells. A full, detailed protocol of tissue dissociation for single cell RNA sequencing can be found at http://www.jasonspencelab.com/protocols.

#### Single nuclei RNA sequencing dissociation

Nuclei were isolated and permeabilized in accordance with 10x Genomics’ Chromium Nuclei Isolation Kit Protocol (10x Genomics Cat#1000493). Briefly, tissue was minced into smaller fragments and then placed in lysis buffer where it was further dissociated mechanically with a pellet pestle. Tissue was then incubated in the lysis buffer for 5-7 minutes. The suspension was passed through the nuclei isolation column and spun at 16,000g for 20 seconds at 4°C. The suspension was then vortexed for 10 seconds and centrifuged at 500g for 3 minutes at 4°C. The supernatant was removed, and the pellet was resuspended in 500 µL of Debris Removal Solution and centrifuged at 700g for 10 minutes at 4°C. The supernatant was removed, and the pellet was resuspended in 1 mL of Wash Solution and centrifuged at 500g for 5 minutes at 4°C twice. The final pellet was resuspended in diluted nuclei buffer. Nuclei capture was carried out on the 10X Chromium platform with a target capture of 5000 nuclei per sample, and libraries were immediately prepared by the University of Michigan Advanced Genomics Core facility.

### Bioinformatics Analysis

#### Sequencing library preparation and transcriptome alignment

All single-cell RNA-seq sample libraries were prepared with the 10X Chromium Controller using v3 chemistry (10X Genomics Cat# 1000268). Sequencing was performed on a NovaSeq 6000 with targeted depth of 100,000 reads per cell. Default alignment parameters were used to align reads to the pre-prepared human reference genome (hg38) provided by the 10X Cell Ranger pipeline. Initial cell demultiplexing and gene quantification were also performed using the default 10X Cell Ranger pipeline.

#### Sequencing data analysis

To generate cell-by-gene matrices, raw data was processed using the 10X Cell Ranger package, and sequenced reads were aligned to the human genome hg38. All downstream analysis was carried out using Scanpy^37^ or Seurat^27^ (depending on package usage needs). For primary human tissue sample analysis in Extended Data Fig. 1, we reanalyzed the human whole cell fetal dataset published in our lab’s previous work^10,19,26^. Samples included a 47-day proximal intestine, a 59-day proximal intestine, two 72-day duodenum, 80-day duodenum and ileum, an 85-day duodenum, 101-day duodenum and ileum, two 127-day duodenums, 132-day duodenum. All samples were filtered to remove cells with less than 500 or greater than 10,000 genes, or greater than 60,000 unique molecular identifier (UMI) counts per cell. De-noised data matrix read counts per gene were log normalized prior to analysis. After log normalization, highly variable genes were identified and extracted, and batch correction was performed using the BBKNN algorithm. The normalized expression levels then underwent linear regression to remove effects of total reads per cell and cell cycle genes, followed by a z-transformation.

Dimension reduction was performed using principal component analysis (PCA) and then uniform manifold approximation and projection (UMAP) on the top 16 principal components (PCs) and 30 nearest neighbors for visualization on 2 dimensions. Clusters of cells within the data were calculated using the Louvain algorithm within Scanpy with a resolution of 1.09. Cell lineages were identified using canonically expressed genes covering 47,100 intestinal cells from all samples.

For Extended Data Fig. 3, all organoid whole cell samples (1 ng/ml EREG, 10 ng/ml EREG, 100 ng/ml EREG, 100 ng/ml EGF) were filtered to remove cells with less than 700 or greater than 6,800 genes, or greater than 33,000 UMI counts per cell, and 0.1 mitochondrial cell counts. Data matrix read counts per gene were log normalized prior to analysis. After log normalization, highly variable genes were identified and extracted, no batch correction was needed as these samples were processed at the same time. Data was then scaled by z-transformation. Dimension reduction was performed using PCA and then UMAP on the top 10 PCs and 15 nearest neighbors for visualization. Clusters of cells within the data were calculated using the Louvain algorithm within Scanpy with a resolution of 0.4. The 10 ng/mL EREG sample alone was filtered to remove cells with less than 700 or greater than 8,000 genes, or greater than 50,000 UMI counts per cell, and 0.1 mitochondrial cell counts. Data matrix read counts per gene were log normalized prior to analysis. After log normalization, highly variable genes were identified and extracted. Data was then scaled by z-transformation. Dimension reduction was performed using PCA and then UMAP on the top 10 PCs and 15 nearest neighbors for visualization. Clusters of cells within the data were calculated using the Louvain algorithm within Scanpy with a resolution of 0.4. The 1 ng/mL EREG sample alone was filtered to remove cells with less than 500 or greater than 8,000 genes, or greater than 45,000 UMI counts per cell, and 0.1 mitochondrial cell counts. Data matrix read counts per gene were log normalized prior to analysis. After log normalization, highly variable genes were identified and extracted. Data was then scaled by z-transformation. Dimension reduction was performed using PCA and then UMAP on the top 10 PCs and 15 nearest neighbors for visualization. Clusters of cells within the data were calculated using the Louvain algorithm within Scanpy with a resolution of 0.4.

For Figure 1, 10 ng/ml EREG single nuclei dataset from *in vitro* grown HIOs was first processed through the standard CellBender^38^ workflow to remove ambient RNA introduced in the nuclei isolation preparation. Then the same dataset was filtered to remove cells with less than 1200 or greater than 6,000 genes, or greater than 17,500 UMI counts per cell, and 0.1 mitochondrial cell counts. Data matrix read counts per gene were log normalized prior to analysis. After log normalization, highly variable genes were identified and extracted. Data was then scaled by z-transformation. Dimension reduction was performed using PCA and then UMAP on the top 18 PCs and 15 nearest neighbors for visualization. Clusters of cells within the data were calculated using the Louvain algorithm within Scanpy with a resolution of 0.5. For Figure 2, the 10 ng/mL EREG single nuclei dataset from transplanted HIOs and was first processed through the standard CellBender^38^ workflow to remove ambient RNA introduced in the nuclei isolation preparation. Then the same dataset was filtered to remove cells with less than 400 or greater than 8,000 genes, or greater than 30,000 UMI counts per cell, and 0.2 mitochondrial cell counts. Data matrix read counts per gene were log normalized prior to analysis. After log normalization, highly variable genes were identified and extracted. Data was then scaled by z-transformation. Dimension reduction was performed using PCA and then UMAP on the top 15 PCs and 15 nearest neighbors for visualization. Clusters of cells within the data were calculated using the Louvain algorithm within Scanpy with a resolution of 0.5.

#### Label transfer of tHIO dataset onto human fetal dataset

We utilized Seurat’s recommended pipeline to perform single-cell reference mapping using the same cells as the reference data (human fetal intestine) and query data (tHIOs). PCAs are first performed on reference and query data. Then a set of anchors are identified and filtered based on the default setting of the function FindTransferAnchors. With the computed anchors, reference.reduction parameter set to PCA, and reduction.model set to UMAP, the function MapQuery returns the projected UMAP coordinates of the query cells mapped onto the reference UMAP. We then integrated the projected UMAP (colored in red) and the reference UMAP (colored in light gray) to visualize the result of our reference-based mapping in Fig. 3.

#### Endoderm atlas

Reference map embedding to the Human Fetal Endoderm Atlas^19^ to determine off target lineages was performed using the scoreHIO R Package. tHIO samples were processed following the preprocessing steps outlined above in the *Single-cell data analysis* section and then put through the basic workflow outlined for this package to map tHIO cells onto the reference endodermal organ atlas.

#### Cell scoring analysis

Cells were scored based on expression of the 20 most differentially expressed genes per tissue type in the human fetal reference dataset. See supplement for gene lists. After obtaining the log-normalized and scaled expression values for the data set, scores for each cell were calculated as the average *z* score within each set of selected genes.

#### tHIO muscle contractions and ENS function

Muscle contraction and ENS function was assayed as previously described^12,39^. Following HIO transplantation as outlined in the previous section, tHIOs were matured for 10-12 weeks before harvest. tHIOs were cut into strips ∼2 × 6 mm in size and the epithelium mechanically removed as previously described^12^. No chelation buffer was used, and all manipulations occurred in oxygenated Kreb’s buffer while on ice ((NaCl, 117 mM; KCl, 4.7 mM; MgCl2, 1.2mM; NaH2PO4, 1.2 mM; NaHCO3, 25 mM; CaCl2, 2.5 mM and glucose, 11 mM), warmed at 37 °C and gassed with 95% O2 + 5% CO2). These strips were mounted in an organ bath chamber system (Radnoti) to isometric force transducers (ADInstruments) and contractile activity was continuously monitored and recorded using LabChart software (ADInstruments). All measurements were normalized to muscle strip mass. After an equilibrium period, a logarithmic dose response to Bethanechol (Sigma-Aldrich Cat#C5259) was obtained through the administration of exponential doses with concentrations of 1 nM to 10 mM at 2 min intervals before the administration of 10 μM scopolamine (Tocris Bioscience Cat#1414/1G). After another equilibrium period, tissue strips were then stimulated with dimethyl phenyl piperazinium (DMPP) (10 μM, Sigma, Cat#D5891). NG-nitro-L-arginine methyl ester (L-NAME) (50 μM, Sigma Cat#N5751) was added 10 minutes before DMPP stimulation to observe the effects of NOS inhibition. Without washing, atropine sulfate salt monohydrate (Atropine) (1 μM, Sigma Cat#A0132) was then applied 10 minutes prior to a final DMPP stimulation to observe the cumulative effect of NOS and Ach receptor inhibition. After several washes and an additional equilibrium period, another dose of DMPP was administered. Neurotoxin tetrodotoxin (TTX) (4 μM, Tocris Cat#1078) was added 5 minutes before a final DMPP stimulation and measurement. Analysis was performed by calculating the integral (expressed as area under the curve, AUC) immediately before and after stimulation for 60 seconds.

### RVEC Experiments

#### Culturing and maintenance

RVECs were obtained from Dr. Shahin Rafii’s Laboratory at Weill Cornell Medicine and generated as previously described^29^. Briefly, various multiplicity of infection (MOI) (from 5 to 20) of lentiviral vectors expressing the transcription factor ETV2 was transduced into human umbilical vein endothelial cells (HUVECs) to generate R-VECs. Then the transduced ECs that generated the most functional perfusable and durable vascular network on the microfluidic devices were selected for further experimentation. We implemented the following protocol to propagate these cells: RVECs were grown in T75 flasks coated in 0.2% gelatin in Endothelial Cell (EC) medium which is comprised of 400 ml M199 (Gibco Cat#11150067), 100 ml HyClone dFBS (Fisher Cat#SH3007103), 5 mL GlutaMAX (1X, Gibco Cat#35050061), 5 mL penicillin-streptomycin (Gibco Cat# 15070063), 7.5 mL HEPES buffer (15 mM, Gibco Cat#15630080), Heparin (Sigma Cat#H3149-100KU), FGF2 (10 ng/mL, R&D Cat#233-FB-MTO), IGF1 (10 ng/mL, Preprotech Cat#100-11), EGF (10 ng/mL, R&D Systems Cat#236-EG-01M) and *N*-acetylcysteine (1.5 mM, Sigma Cat#A9165-25G). The cells were split 1:3 using Accutase (Corning Cat#MT25058CI) and passaged on gelatin coated flasks.

#### Lentiviral labeling

RVECs were transduced with GFP lenti-particles (Lenti-EV-GFP-VSVG) provided by the University of Michigan Vector Core. Virus was diluted in EC media and added to the RVECs for 8 hours. After RVECs were incubated with the virus, the cells were thoroughly washed and allowed to continue to grow normally.

#### Microfluidic device

Polydimethylsiloxane (PDMS; Sylgard 184; Ellsworth Adhesives Cat#2065622) based microfluidic devices were fabricated via soft lithography with a 3D printed resin cast. The device is 50mm in length, 20.64mm in width, and 3mm in height. The physical chamber housing the 3D co-culture carries a height of 1.5mm to account for larger size HIOs. Each device was plasma treated, with Harricks Expanded Plasma Cleaner (Harricks Plasma Cat#PDC-001), to a 24×60mm glass cover slip (VWR Cat#152460), and then placed in an 80°C oven for at least 1 hour to finalize a strong adhesion. For long term storage, devices were sealed with parafilm. Before use, devices were sterilized with UV light for at least 30 minutes prior to seeding of cells.

RVECs were washed with sterile PBS then incubated in accutase for 3-5 minutes. Digestion was stopped by adding an equal volume of EC media and cell suspension was obtained by centrifugation at 500g for 5 minutes at 4°C. Supernatant was removed and RVECs were resuspended in M199 (Gibco Cat#11150067) and counted. 250,000 RVEC was aliquoted into a 1.5mL micro-centrifuge tube, which corresponds to a single lane on the microfluidic device. HIOs were removed from Matrigel using a cut P200 tip and transferred to a 1.5mL micro-centrifuge tube to be spun at 300g for 5 minutes at 4°C. The supernatant was removed and resuspended in DMEM. Three to five HIOs were then picked and added to each aliquot of RVECs in the 1.5mL micro-centrifuge tube, which was subsequently centrifuged at 500g for 5 minutes at 4°C. Supernatant was removed and resuspended in 32uL of a Fibrin mixture consisting of Fibrinogen from bovine plasma (Sigma Cat#F8630), Human Fibrinogen 1 Plasminogen Depleted (Enzyme Research Lab Cat#FIB-1), and X-Vivo 20 (Lonza Cat#190995). 3.6uL of a Thrombin mixture, consisting of Thrombin from bovine plasma (Sigma Cat#T4648) and X-Vivo 20, was then added to the mixture of RVECs and Fibrinogen. The final matrix concentration is 0.5mg/mL of Human Fibrinogen, 2mg/mL of Bovine Fibrinogen, and 2U/mL of Thrombin. The cell mixture was resuspended and immediately seeded into the microfluidic chamber within 10-15 seconds. Between loading each lane, devices were flipped upside down to prevent HIOs resting to the bottom. Devices were then incubated, right side up, for 5-15 minutes.

40uL of Tube Media (500mL StemSpan SFEM, Stemcell Technologies Cat#9650; 50mL Knockout Serum, Gibco Cat#10828010; 5mL Penicillin-Streptomycin, Gibco Cat#15070063; 5mL Heparin, Sigma Cat#H3149; 5mL GlutaMAX, 1X Gibco Cat#15070063; 5mL HEPES Buffer, Gibco Cat#15630080; 10ng/mL FGF2, R&D Cat#233-FB-MTO; 10ng/mL Aprotinin, Sigma Cat#A6106) was added to both inlet and outlet. A 1mL syringe, without the plunger, was additionally attached to both ends and 1mL of Tube media was added to the inlet syringe to induce shear stress via gravity. Media in the outlet was recycled back to the inlet on a daily basis.

## Data and code availability statement

Sequencing data generated and used by this study are deposited at EMBL-EBI ArrayExpress. Data sets for human fetal intestine (ArrayExpress: E-MTAB-9489, https://www.ebi.ac.uk/arrayexpress/experiments/E-MTAB-9489/, and previously published work^26^); Data sets for whole cell single cell sequencing of 1 ng/mL, 10 ng/mL, 100 ng/mL EREG HIOs and 100 ng/mL EGF HIO (ArrayExpress: E-MTAB-13463, https://www.ebi.ac.uk/arrayexpress/experiments/ E-MTAB-13463/,); Data sets for single nuclear RNA sequencing of 10 ng/mL HIOs and 10 ng/mL EREG tHIOs (ArrayExpress: E-MTAB-13469, https://www.ebi.ac.uk/arrayexpress/experiments/ E-MTAB-13469/). Code used to process raw data can be found at https://github.com/jason-spence-lab/Childs_2023.git

---

**Extended Data Figure 1:**
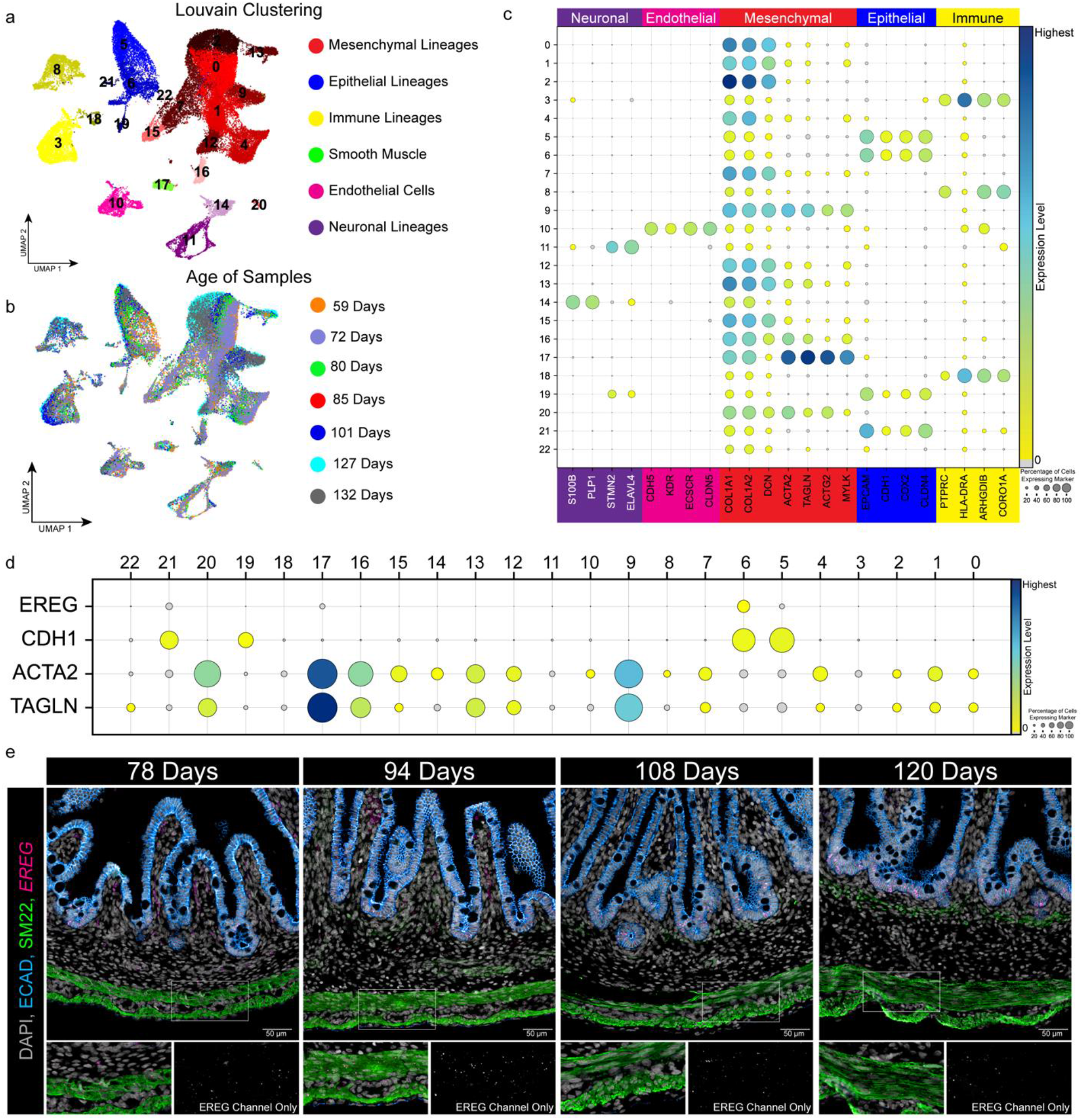
EREG is expressed in the developing human intestine’s crypts and smooth muscle bands across developmental time. a) UMAP visualization of Louvain clustering by cell type of human fetal intestine (51,790 cells, n=13 biological samples) for major tissue classifications b) UMAP visualization of clustering by age of human fetal intestine (n=13 biological samples. 59-, 72-, 80-, 85-, 101-, 127-, and 132-days post conception). c) Dot plot of fetal dataset highlighting expression of canonical lineage genes used for cluster annotation by tissue type. d) Dot plot of EREG expression in epithelial clusters (marked by CDH1) and smooth muscle clusters (marked by ACTA2 and TAGLN). e) Co-FISH/IF staining for *EREG* (pink), DAPI (grey), ECAD (blue), and SM22 (green) in the developing human intestine at select timepoints across developmental time.

**Extended Data Figure 2:**
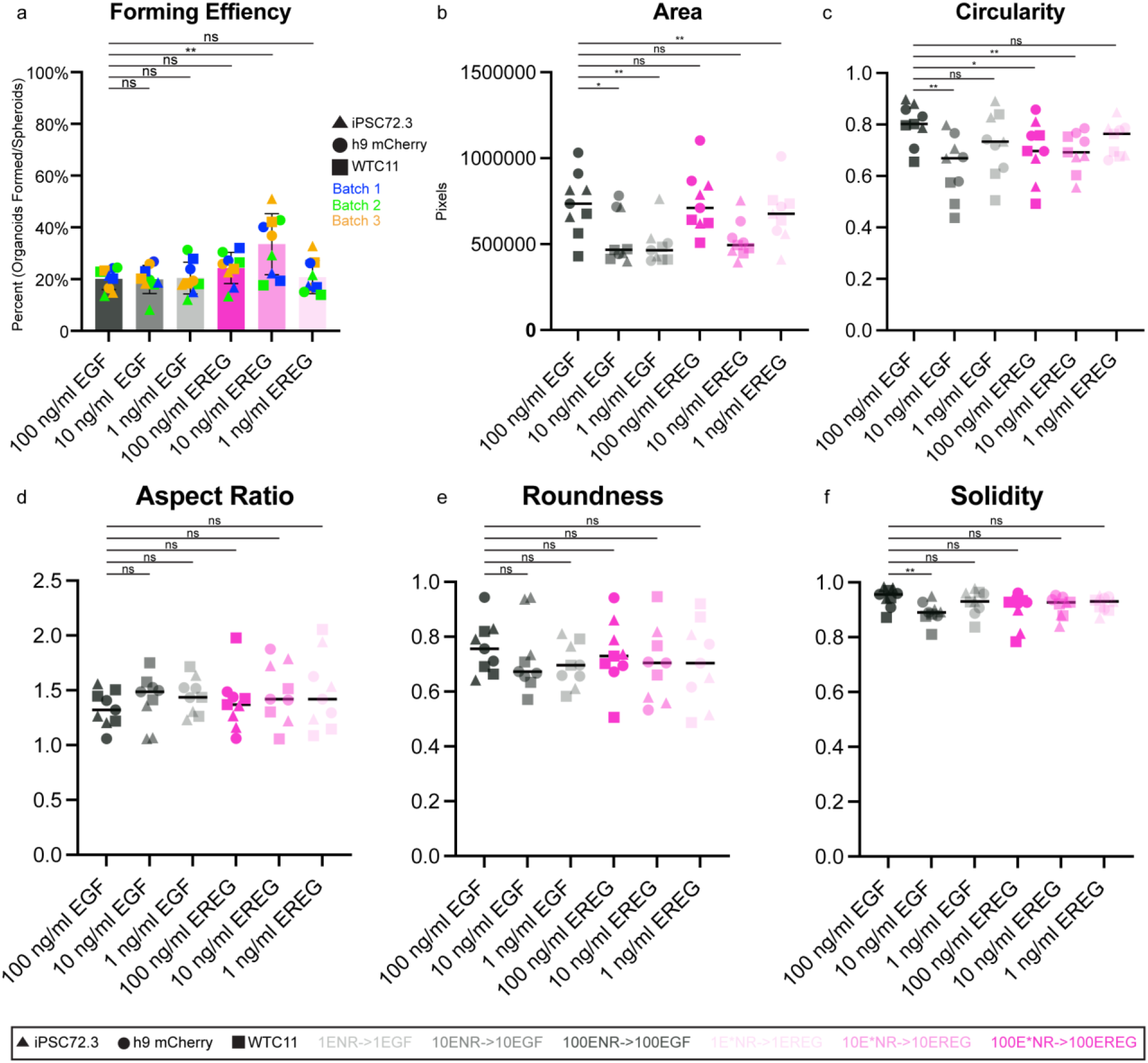
Measurements of forming efficiency, area, and shape of control EGF-grown HIOs compared to EREG-grown HIOs. a) Organoid forming efficiency assay results (n = 3 biological replicates with n = 3 technical replicates quantified for three different cell lines). Statistical significance was determined using an unpaired Welch’s t test (** - P = 0.0097, ns – P = >0.05). b-f) Morphologic quantifications including area, solidity, aspect ratio, circularity, and roundness of HIOs grown in varying doses of EGF or EREG. HIOs were derived from three separate cell lines and grown for 30 days. Three HIOs per condition were measured and the ImageJ analysis software was used to calculate these measurements. See methods section for further explanation on calculations. Statistical significance was determined using an unpaired Welch’s t test (ns – P = >0.05).

**Extended Data Figure 3:**
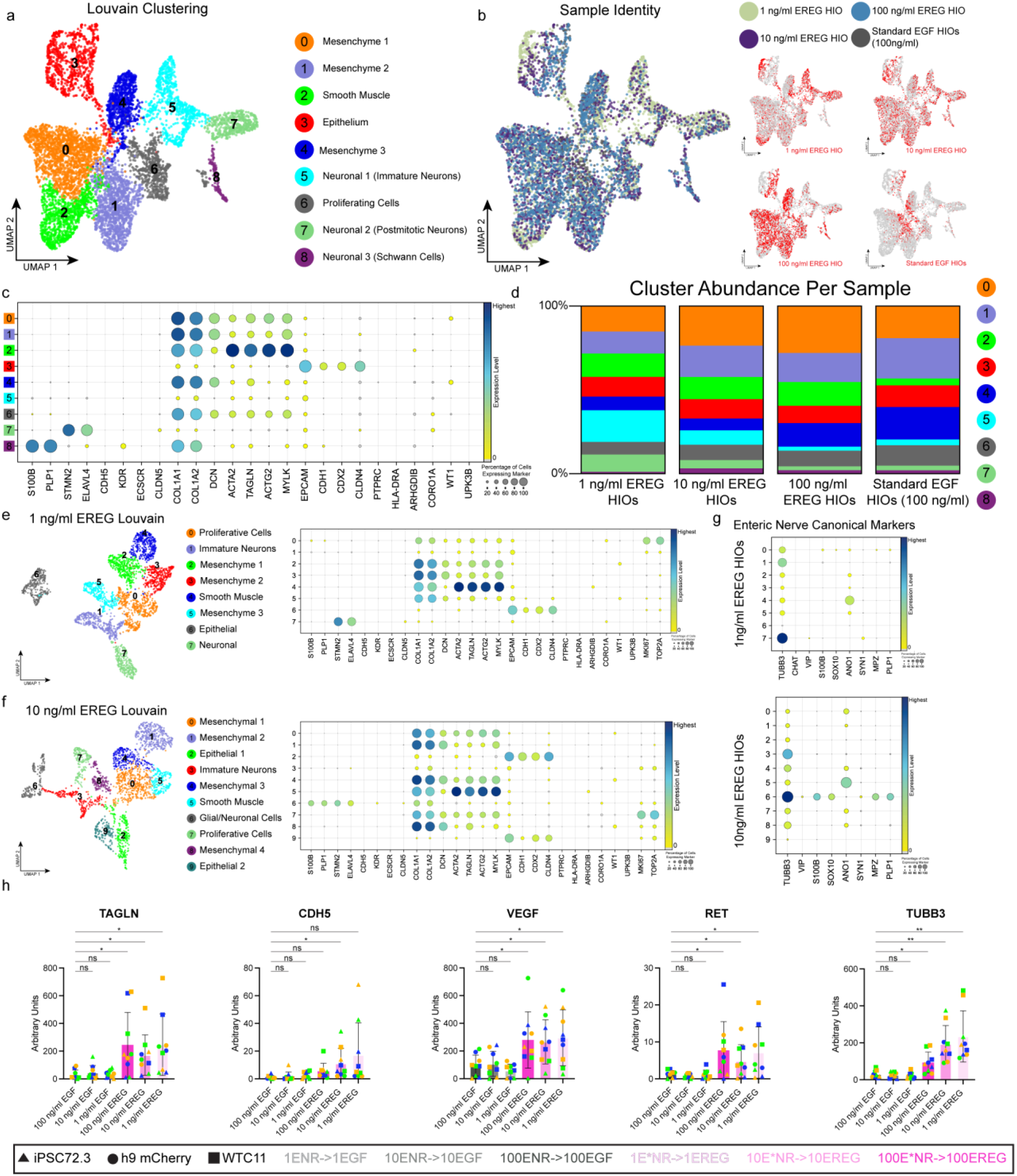
Gene expression analysis of EREG-grown HIOs show presence of smooth muscle, neural cells, and endothelial cells not seen in control EGF-grown HIOs. a) UMAP visualization of Louvain clustering by cell type of all HIO samples sequenced (100 ng/mL EREG, 10 ng/mL EREG, 1 ng/mL EREG, and 100 ng/mL EGF). b) UMAP visualization of clustering by sample type (100 ng/mL EREG blue, 10 ng/mL EREG purple, 1 ng/mL EREG green, and 100 ng/mL EGF grey). Inlays break out each sample (red) individually against rest of dataset (grey). c) Dot plot of combined HIO dataset highlighting expression of canonical lineage genes used for cluster annotation. d) Bar charts showing the cell type abundance (% of total cells) within each cluster for each sample sequenced. Colors are consistent with the cell type annotation in panel A. e) UMAP visualization of Louvain clustering by cell type of 1 ng/mL EREG and accompanying dot plot of expression of canonical lineage genes used for cluster annotation. f) UMAP visualization of Louvain clustering by cell type of 10 ng/mL EREG and accompanying dot plot of expression of canonical lineage genes used for cluster annotation. g) Dot plot of individual HIO dataset highlighting expression of major ENS neuronal cell types seen in the developing human intestine. Enteric ganglion cells (*TUBB3, SYN1*), submucosal secretomotor (*VIP*), enteric glial cells (*S100b* - glial network; *SOX10* - EGC nuclei), ICC’s (*ANO1*), cholinergic neurons (*CHAT*) and Schwann cells (*MPZ, PLP1*). h) Bar charts showing gene expression of smooth muscle (*TAGLN*), endothelial cells (*CDH5, VEGF*) and Neurons (*RET, TUBB3*) for 100 ng/mL, 10 ng/mL, and 1 ng/mL EREG and matched EGF HIOs. Data points shown are the average of triplicates completed in 3 different passages (batches) for 3 different cell lines. Statistical significance was determined using an unpaired Welch’s t test to the standard 100 ng/mL EGF condition (ns – P = >0.05).

**Extended Data Figure 4:**
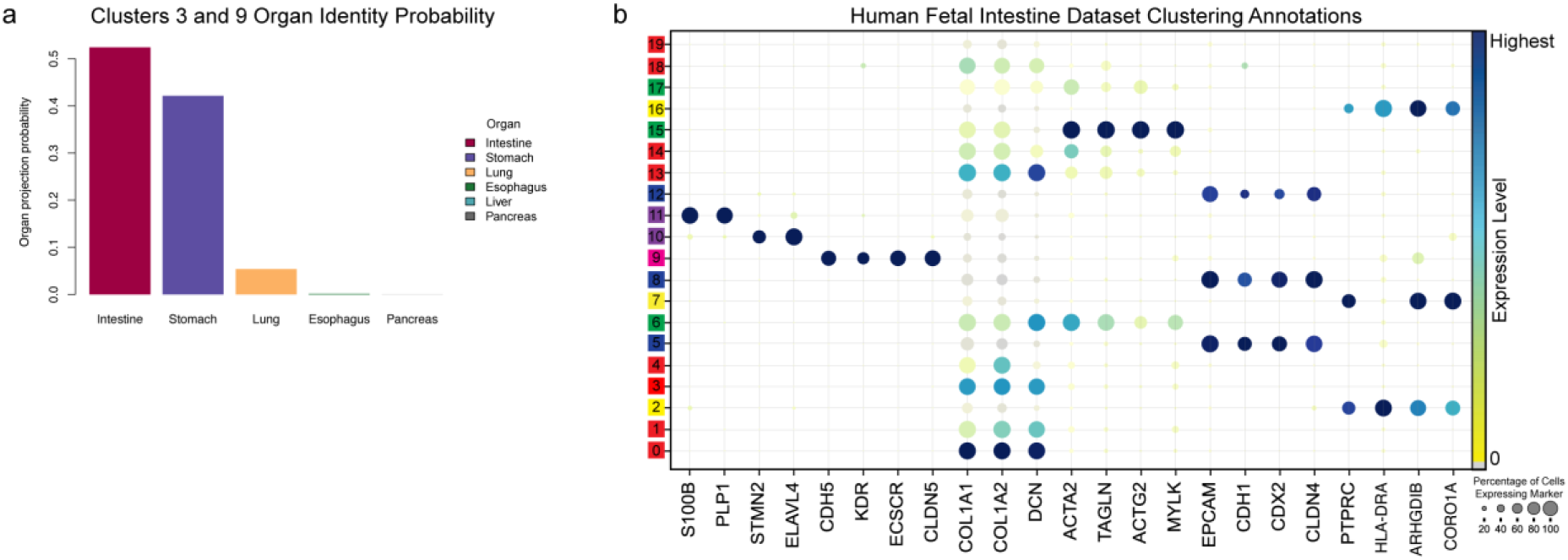
EREG-grown tHIO pattern two off target epithelial clusters. a) Bar plot of clusters 5 and 7 showing predicted organ identity using the scoreHIO R package. Clustered mapped somewhere between gastric epithelium and intestinal epithelium. b) Dot plot of human fetal dataset highlighting expression of canonical lineage genes used for cluster annotation in label transfer and SingleR analysis.

**Extended Data Figure 5:**
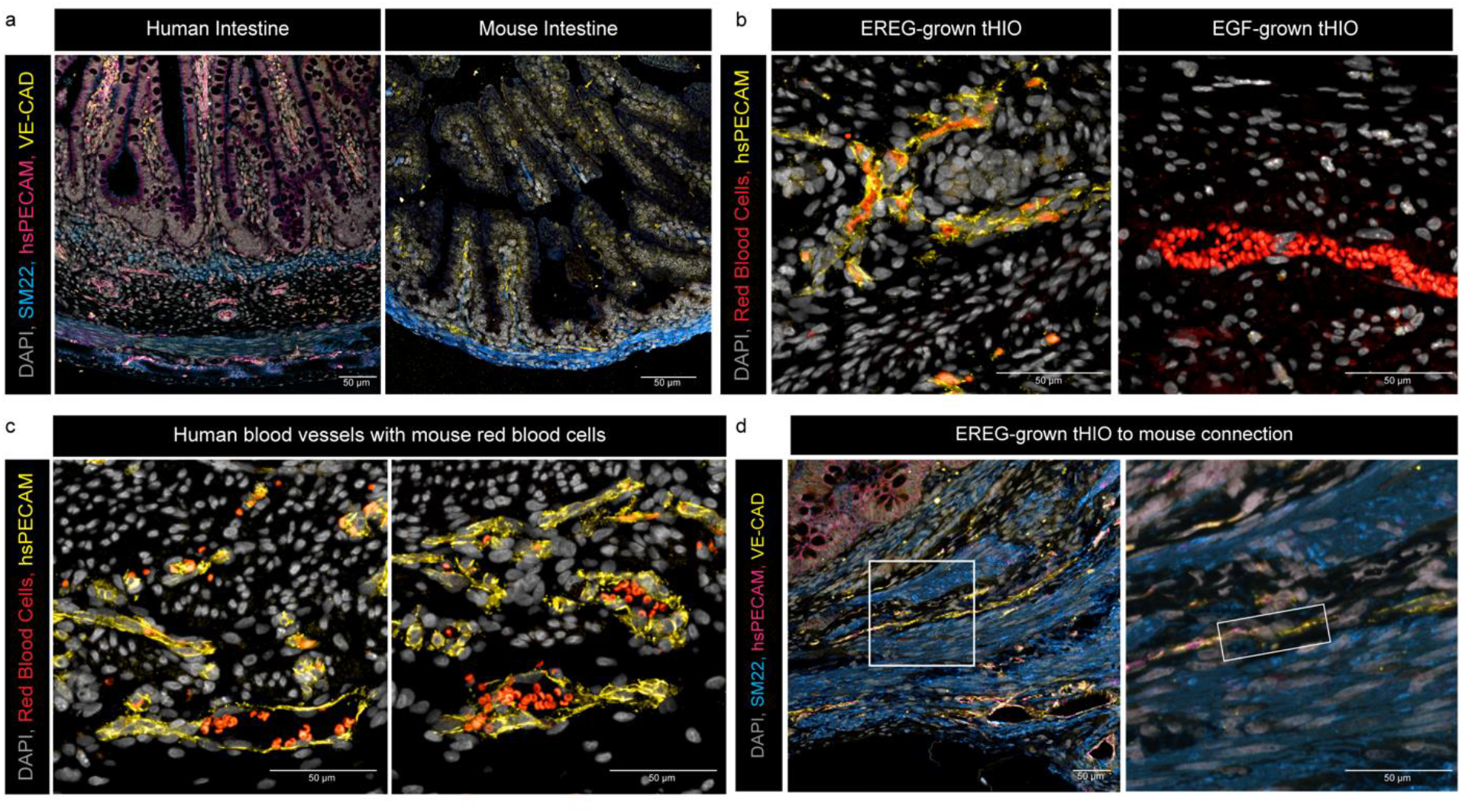
EREG-grown tHIOs feature human blood vessels that can connect to their murine host. a) IF staining of human fetal intestine (Left; 127-days post conception) and E13.5 mouse intestine to test for human specificity of hsPECAM (pink) antibody compared to a pan-species VE-CAD antibody (yellow) and smooth muscle SM22 (blue). All Scale bars = 50 µm. b) IF staining of EREG-grown tHIO (Left) and EGF-grown tHIO (Right) with stains for DAPI (grey), autofluorescent red blood cells in the 488-laser channel (red), and human specific PECAM antibody (yellow). All scale bars = 50 µm. c) IF staining of other areas of EREG-grown tHIO with stains for DAPI (grey), autofluorescent red blood cells in the 488-laser channel (red), and human specific PECAM antibody (yellow). All Scale bars = 50 µm. d) IF staining of EREG-grown tHIO vessels (co-stain of hsPECAM in pink with pan-species VE-CAD in yellow) connecting to a mouse blood vessel (stained for only pan-species VE-CAD yellow) with smooth muscle SM22 (blue). All Scale bars = 50 µm.

**Extended Data Figure 6:**
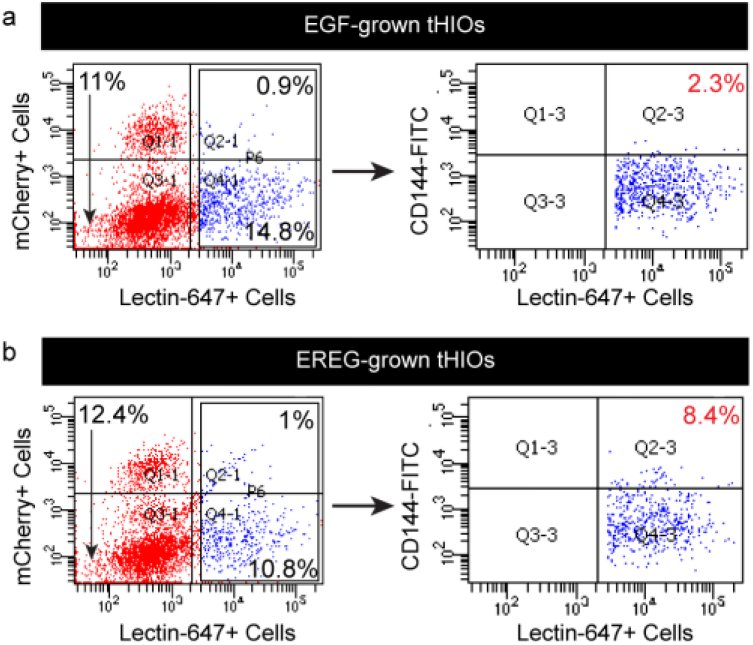
Flow plots for human specific lectin vessel quantification. a) Flow cytometric plots of batch-matched 12-week control EGF-grown HIOs for mCherry tag, human specific CD144/VE-CAD, and Lectin. Flow cytometric analysis required multiple tHIOs to be pooled from the same batch to ensure enough cells for experiment. b) Flow cytometric plots of batch-matched 12-week EREG-grown HIOs for mCherry tag, human specific CD144/VE-CAD, and Lectin. Flow cytometric analysis required multiple tHIOs to be pooled from the same batch to ensure enough cells for experiment.

## Notes

### Competing Interest Statement

Charlie J. Childs, Emily M. Holloway and Jason R. Spence hold intellectual property pertaining to human intestinal organoids.

### Summary of Updates

Addition of author's middle initial and ORCID

